# Understanding the contributions of visual stimuli to contextual fear conditioning: a proof-of-concept study using LCD screens

**DOI:** 10.1101/069781

**Authors:** Nathen J. Murawski, Arun Asok

## Abstract

The precise contribution of visual information to contextual fear-learning and discrimination has remained elusive. To better understand this contribution, we coupled the context pre-exposure facilitation effect (CPFE) fear conditioning paradigm with presentations of distinct visual scenes displayed on 4 LCD screens surrounding a conditioning chamber. Adult male Long-Evans rats received non-reinforced context pre-exposure on Day 1, an immediate 1.5 mA foot shock on Day 2, and a non-reinforced context test on Day 3. Rats were pre-exposed to either digital Context (dCtx) A, dCtx B, a distinct Context C, or no context on Day 1. Context A and B were identical except for the visual image displayed on the LCD monitors. Immediate shock and retention testing occurred in dCtx A. Rats pre-exposed dCtx A showed the CPFE with significantly higher levels of freezing compared to learning controls. Rats pre-exposed to Context B failed to show the CPFE, with freezing that did not differ significantly from any group. The results suggest that 1) visual information contributes to contextual fear learning in rats and that 2) visual components of the context can be parametrically controlled via LCD screens. Our approach offers a simple modification to contextual fear conditioning whereby the visual features of a context can be precisely controlled to better understand how rodents discriminate and generalize fear across environments.

## 1. Introduction

In fear conditioning paradigms, manipulating contextual cues is important for demonstrating that learning is either ^“^context-specific^”^ or ^“^context-independent^”^ given that distinct (although possibly overlapping; (Cai et al., 2016; Rashid et al., 2016)) neural systems support these different types of conditioned fear (Fendt & Fanselow, 1999; LeDoux, 2000). ^“^Context,^”^ in fear conditioning, is defined as the multimodal sensory experience, including temporal and spatial factors, encountered concurrently during a conditioning trial (for reviews see (Holland & Bouton, 1999; Rudy, 2009; Smith & Bulkin, 2014). In contextual fear conditioning, the context is usually the conditioning chamber, an enclosed box with visual, tactile, olfactory, and auditory cues that are incidentally encountered and associated with an aversive stimulus (e.g., foot shock). Following a conditioning trial, a rodent will typically freeze (a species-specific specific defensive reaction; (Blanchard & Blanchard, 1969; Bolles, 1970)) when placed back into the context where it previously encountered the foot shock, but will not freeze if placed into a novel context (i.e., the rodent’s behavioral response is context-specific). However, it is unclear as to what makes one context distinct from another. Typically, researchers manipulate multiple sensory features of a context to make one context distinct from another. Yet, our understanding of how the independent sensory features of a context may differentially contribute to the CS-US representation (i.e., is one feature more prominent than another?) and what specific elements are most important in making one context distinct from another is limited given that researchers often use gross manipulations of contextual cues (Fanselow, 2000).

Rodents are able to utilize visual information presented on LCD screens to perform a variety of behavioral tasks. For example, rodents can successfully traverse a virtual maze or use digital images to make appropriate behavioral choices (Lee & Shin, 2012; Swan et al., 2014; Thomas et al., 2007). In addition, presenting looming or sweeping visual stimuli – to simulate predator approaching or cruising behaviors - on a ceiling-mounted LCD screen can control rodent flight or freezing behavior, respectively (De Franceschi et al., 2016). Incorporating LCD monitors in a contextual fear conditioning paradigm may offer a means to systematically control visual features of a context during aversive learning.

To examine this possibility, we utilized a variant of contextual fear conditioning known as the context pre-exposure facilitation effect (CPFE) paradigm. The CPFE paradigm separates incidental contextual learning from context-shock associative learning. The CPFE relies on the immediate shock deficit (ISD) – a phenomenon where animals that are not given enough time to learn about the context prior to receiving a shock (e.g., < 10-sec (Fanselow, 1986)) fail to exhibit conditioned freezing (Burman, Murawski, Schiffino, Rosen, & Stanton, 2009; Fanselow, 1990)). However, pre-exposure to the conditioning context on the day prior to receiving an immediate shock is sufficient to overcome this deficit (Fanselow, 1990). More importantly, the CPFE is only evident when context pre-exposure occurs in the conditioning context and not a distinct context (Rudy, Huff, & Matus-Amat, 2004). While the CPFE paradigm has been useful in understanding how gross incidental contextual learning is processed distinctly from fear-learning (Matus-Amat, Higgins, Barrientos, & Rudy, 2004; Matus-Amat, Higgins, Sprunger, Wright-Hardesty, & Rudy, 2007), the quantitative contribution of visual features to contextual learning remains unclear.

In the present study, we investigated how altering the visual environment affects contextual fear-learning. We restricted changes in the visual environment to incidental contextual learning during context pre-exposure. We placed four LCD monitors around a clear chamber (three on the sides and one on top). The LCD monitors displayed one of two images on all screens during the pre-exposure phase, with all other context features (i.e., tactile, spatial, olfactory, and auditory components) held constant between the two groups. During immediate-shock training and testing, only one of the two visual scenes was displayed on the monitors. We hypothesized that that rats pre-exposed to the testing context would show the CPFE whereas those pre-exposed to the alternate visual context would not. We also included two control groups, one of which was pre-exposed to a distinct context (different visual, spatial, auditory, and olfactory components) and one group that received no pre-exposure as CS and US associative-learning controls.

## 2. Methods

### 2.1. Subjects

Forty adult male Long Evans rats 8-9 weeks of age were used in the present study. Thirty-two were purchased from Harlan breeders (Indianapolis, IN) and eight were bred in house (University of Delaware). All rats were housed in the animal colony at the University of Delaware. Pairs of rats were housed in opaque polypropylene cages (45×24×21 cm) with standard bedding and free access to food and water. Rats were maintained on a 12:12 h light/dark cycle with lights on at 7:00 am. All behavioral testing occurred during the light phase between (12:00PM – 5:00PM). All animals were treated in accordance with NIH guidelines for the care and use of laboratory animals.

### 2.2. Apparatus

The conditioning chamber was made out of clear Plexiglas (40x22x24 cm); one of the four walls could be opened to allow for animal placement. The chamber floor consisted of 40 grid bars (0.4 cm in diameter) that ran parallel to the shorter wall of the chamber. The grid bars were connected to a shock generator (Med Associates; ENV 410B) that delivered an alternating current foot shock US. Four LCD monitors (Dell, Plano, TX) were placed flush to three of the external walls of the chamber with one monitor acted as the ceiling. All monitors projected the same image concurrently and were connected to a Dell computer. The monitors projected one of two images. The images used were found on an internet search and did not have copyright attribution. The first image consisted of multiple pumpkins with dominant ovoid contours and light and dark coloring throughout. The second image consisted of rock formations with dominant linear contours and darker coloring toward one side, lighter coloring towards the other side. The lighting levels between the two images were adjusted via the LCD monitors so that their total luminance (∼200-225 lux) was equivalent. Image presentation was controlled by custom software (available at: https://sites.google.com/site/aaasok/programmed-software). The conditioning chamber was cleaned with a 70% ethanol solution prior to animal placement. When the monitors projected the pumpkin image, the chamber was determined to be in the Context A configuration; When the monitors projected the rock formation image, the chamber was determined to be in the Context B configuration (Figure 1A). With the exception of the projected images, there were no other differences between Context A and B.

**Fig. 1.**
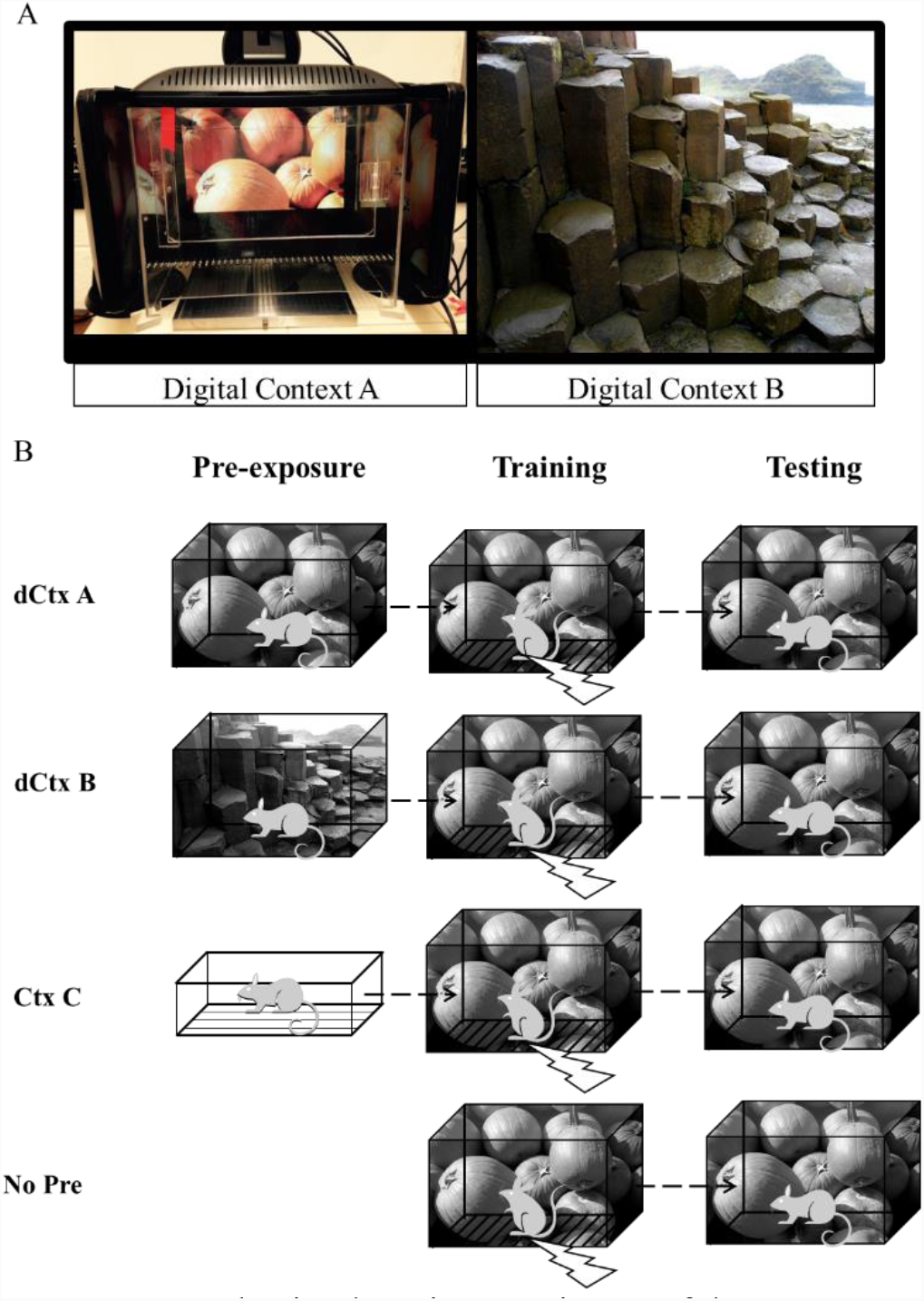
CPFE Behavioral Design. (A) image of the conditioning apparatus with LCD screens and digital context (dCtx) A (left panel) and dCtx B (right panel). (B) The CPFE fear conditioning experiments were run over three days. On day 1, rats were pre-exposed to digital context (dCtx) A, dCtx B, Ctx C, or remained in their homecages with no pre-exposure. On day 2, all groups were given immediate shock training in dCtx A. On day 3, all groups were returned to dCtx A and tested for fear conditioned freezing. dCtx A and dCtx B were identical except for either pumpkins or a rock formation displayed on the LCD screens. Ctx C was distinct with regard to spatial, tactile, visual, olfactory, and auditory characteristics. No Pre animals were not pre-exposed to any context.

The conditioning chamber was located in a dark room with low-levels of background noise produced by ventilation. A CCD camera (Model # ACT-VP-02, Actimetrics, Wilmette, IL) mounted on a tri-pod was placed approximately 61 cm in front of the clear wall of the conditioning chamber. The camera was connected to a computer running FreezeFrame3 software (Actimetrics, Wilmette IL), which controlled session protocol (see below) and foot shock presentation.

A separate Plexiglas chamber (16.5 x 21.1 x 21.6 cm) was used as Context C. This chamber was situated in a separate, dark room within a fume hood that provided ambient noise (∼74 db) and lighting (∼1200 lux). The chamber had a grid floor with 9 metal bars (0.5 cm in diameter and placed 1.25 cm apart) parallel to the long length of the chamber. One side of the chamber was painted white. The chamber was cleaned with a 5% ammonium hydroxide solution prior to animal placement.

### 2.3. Procedure

2.3.1. General Procedure

Four groups of rats were included in the experiment. One group received no pre-exposure (group No Pre), one group was pre-exposed to Context C (group Ctx C), one group was pre-exposed to digital context B (group dCtx B), and one group was pre-exposed to digital context A (group dCtx A). Prior to each behavioral session (see below) rats were brought up from the animal colony in their home cages and then placed on a cage rack in a waiting room adjacent to the testing room. Rats were run one at a time. One animal was taken from its home cage and placed into a black ice bucket with the cover on and transferred to the testing room. Each rat was quickly removed from the ice bucket and placed into the conditioning chamber (< 30 sec.). At the end of the session, the rat was quickly removed from the conditioning chamber and placed back into the ice bucket. The rat was then transferred to a clean holding cage while the experimenter cleaned out the conditioning chamber and ice bucket in preparation for the next rat.

### 2.3.2. Behavioral Procedure (Figure 1B)

Figure 1 provides a schematic of the behavioral testing procedure, which included three sessions (Context pre-exposure; Immediate shock training; and Context testing). Each session occurred 24-hours apart.

For *context pre-exposure*, rats were pre-exposed to one of three contexts (digital contexts A or B, or alternate context C). A fourth group was briefly removed and returned to their home cage to parallel handling by the experimenter similar to what was experienced by the other groups. Context pre-exposure consisted of a 300-sec session in the absence of any US presentation. Additionally, to match the tactile experience across digital contexts, a dark grey Plexiglas sheet was placed over the grid floor.

For *immediate shock training*, rats were placed into digital Context A and received an immediate (< 5-sec) 1.5 mA, 2-sec AC foot shock US. Rats were removed from the conditioning chamber within 5-sec of foot shock presentation.

For *context testing*, rats were placed back into digital Context A and their freezing behavior was monitored over a 300-sec testing session. Prior to placement, the dark grey Plexiglas floor covering (that was used during pre-exposure) was placed over the grid floor.

### 2.4. Data Analysis

#### 2.4.1. Freezing Analyses

The data were collected and scored using FreezeFrame3 software (Actimetrics, Wilmette IL). The bout length was set to 0.75-sec and the freezing threshold (changes in pixels per frame) was adjusted by an experimenter so that no small movements were registered as freezing.

#### 2.4.2. Statistical Analyses

SPSS 23.0 was used for all data analyses (IBM Corp, Armonk, NY). Both pre-exposure and testing data were analyzed separately with a repeated-measures ANOVA across five 60-sec time bins as the within-subject’s variable and group (Pre-exposure context) as the between-measures variable. We used this design because (1) the No Pre group was not pre-exposed to any context and thus had no pre-exposure data and (2) we were interested in how freezing changed within a session depending on the visual environment. Post-hoc ANOVA’s and Dunnet’s 2-sided contrasts were used to follow up on significant interactions to examine differences between groups and relative to the associative learning control group (Group No Pre) at specific time bins. Additionally, we used linear regression analyses to ask what the on-average change in freezing was across groups as the context became successively closer to the conditioning context. Significant effects were set at *p* < 0.05.

## 3. Results

3.1. Pre-exposure (Figure 2A)

**Fig 2.**
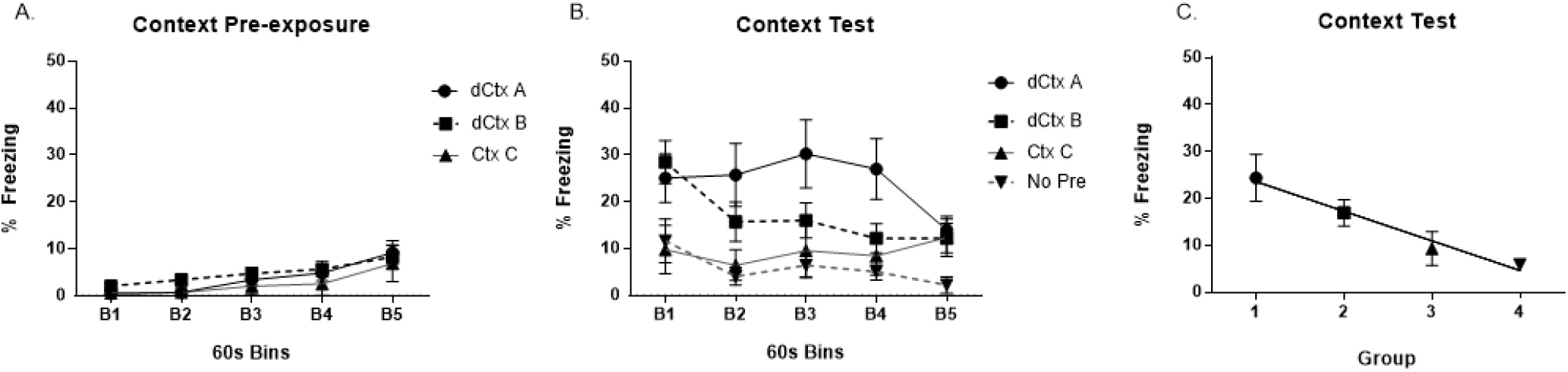
Rats were pre-exposed to digital Context A (dCtx A), digital Context B (dCtx B), or an alternate context (Ctx C). The percentage of time rats froze during the 300 second context pre-exposure session (A). All groups showed low levels of freezing during context pre-exposure with no differences between groups. During the context test, rats pre-exposed to dCtx A showed significantly higher levels of freezing across the 300 second context testing session relative to rats without context pre-exposure (B). Rats pre-exposed to dCtx B did not differ from any group. The average percentage freezing during Context Testing (C). Group 1=dCtx A, Group 2=dCtx B, Group 3=Ctx C, and Group 4=No Pre. A regression line shows the relationship between groups, indicating increased levels of context mediated freezing as the pre-exposure context shared more features to the testing context.

Freezing levels during the pre-exposure session were low and did not differ among groups (freezing less than 3% overall). Only three groups, dCtx A _n= 14_, dCtx B _n = 14_, and Ctx C _n = 6_ received context pre-exposure. The No Pre n = 6 group did not receive context pre-exposure and was excluded from pre-exposure analysis. A significant main effect of bin *F*(1.928, 124)=5.716, *p*< .01 (Greenhouse-Geisser corrected) was detected. However, there was no main effect of group F(2,31)=.428, *p*> .05 and no Group X Bin interaction F_3.856, 124_=.990, *p*>.05 (Greenhouse-Geisser corrected).

### 3.2. Testing (Figure 2B and 2C)

The amount of context-mediated freezing during the testing session differed among pre-exposure groups. During the 5-minute test, analysis revealed a significant main effect of Time bin [F_4_, _144_=3.109, p<0.05], a significant main effect of group [F_3,36_=3.668, *p*<.05], and a significant Time bin x Group interaction [F_12, 144_=1.885, *p*<0.05]. Follow-up ANOVAs showed that groups marginally differed at bin1 [F_3, 36_=2.581 *p*=.069], significantly differed at bin 2 [F_3, 36_=3.186 *p*<.05], bin 3 [F_3, 36_=3.532 *p*<.05], and bin 4 [F_3, 36_=3.799 *p*<.05]. Groups did not significantly differ at bin 5. Post-hoc 2-sided Dunnet’s test revealed that group dCtx A significantly differed from the No Pre group at all significant bins (*p*’s <.05). dCtx B and Ctx C did not significantly differ from group No Pre (*p*’s > .05). A t-test contrasting dCtx A group vs. dCtx B group showed a marginal difference at bin 3 t(26)=2.006, *p*=.055 and significant difference at bin4 t(26)=2.088, *p*<.05.

Additionally, a significant linear regression equation was found [F_1, 38_=11.091, *p*<.01], with an R^2^ of .226. On average, a group’s freezing increased (from 3.21%) by 7.5% as the pre-exposure context became more similar to the conditioning context.

## 4. Discussion

In the present study, we asked if changes to visual information in a context– changes systematically manipulated using LCD screens – during context pre-exposure mediate conditioning to an immediate shock subsequently presented. Rats pre-exposed to the testing context (group dCtx A) demonstrated the CPFE, wherein context pre-exposure facilitated context conditioning to an immediate shock; this facilitation was relative to controls. Both control groups (i.e., groups CtxC and No Pre) displayed the immediate shock deficit, with little context-mediated freezing evident during testing. In contrast, those rats pre-exposed to the testing context with an alternate visual image displayed on the LCD screens (group dCtx B) failed to show the CPFE, with freezing levels that were not significantly different from either group dCtx A or controls. Because rats from groups dCtx A and dCtx B were trained under identical contextual parameters (i.e., olfactory, tactile, visual, auditory, etc.) the differences in context-mediated freezing are attributable to the visual images displayed during the pre-exposure session (i.e., identical contextual parameters except visual features). Additionally, as the pre-exposure context shares more features with the testing context, animals on average increase their freezing by about 7.5%. These data highlight how (1) visual information from an environment may statistically contribute to contextual fear learning and (2) LCD monitors can be readily incorporated to current fear conditioning setups within laboratories in order to precisely control visual stimuli.

During context pre-exposure, rats from each of the three pre-exposure groups displayed low levels of freezing, an indirect measure of context exploration. Towards the end of the pre-exposure session, freezing levels began to increase slightly, but did not differ among groups. This reduction in exploration during the latter portions of the pre-exposure session may reflect habituation to the context, an indirect measure of context learning (Stote & Fanselow, 2004). In the current study, the lack of a difference in freezing during context pre-exposure between groups may indicate that each group learned about their respective pre-exposure contexts to a similar degree.

It is interesting to note that during testing, both visual context groups show similar levels of freezing in the first minute, but begin to differ as the session progresses. During the CPFE, the context memory associated with the immediate shock is that of the pre-exposure context (Rudy, Barrientos, & O’;Reilly, 2002). We speculate that group dCtx B likely formed a contextual fear memory to dCtx B. During the initial moments of the context test in dCtx A, it is possible that this group generalized fear to those features of the testing environment that were shared with the pre-exposure environment (e.g., tactile, spatial, etc.). Over the testing session, however, it is possible that these rats presumably sampled additional features of the testing context (i.e., visual information) that were not shared with the pre-exposure environment thus facilitating context discrimination. Although, we did not test dCtx B rats for freezing in dCtx B which limits the interpretation of what was likely occurring in this group to speculation.

Recent computational models of contextual fear conditioning predict this effect. For example, the contextual fear conditioning model developed by Krasne, Cushman, and Fanselow (2015) relies on the assumption that only a small subset of context features can be sampled at any given time; further, that over time there is a continual re-evaluation of context features to determine if the context as a whole is either familiar (i.e., previously experienced) or novel. This model predicts greater generalization between more familiar contexts. Specifically, those contexts which share more features will initially produce high-levels of fear during testing, but this fear will quickly dissipate due to extinction (Krasne, Cushman, & Fanselow, 2015). Our data lend experimental support to this model. First, when few context features were shared between the pre-exposure and testing environment there was little generalization (i.e., group Ctx. C). However, when all but one (the visual) of the context features were shared between the environments, there was initially high generalization (i.e., more freezing) that quickly decreased across the testing session.

Given that contextual fear conditioning is thought to involve a conditioned stimulus that is a composite product of a multisensory experience (Maren, Phan, & Liberzon, 2013), isolating the unique contributions of each feature in the environment - the tactile experience from the bars, the olfactory stimuli from cleaning agents, the spatial dimensions of the chamber, etc. - has proved challenging. Previous studies have used gross manipulations of one or more sensory features to make one context distinct from another (e.g., (Bucci, Saddoris, & Burwell, 2002; Nakashiba et al., 2012; Wiltgen et al., 2010). Lee and colleagues (Kim, Lee, & Lee, 2012) outline a number of limitations inherent in gross manipulations of context features; these include vaguely defined contextual stimuli and a lack of experimental control over within-session changes to those stimuli. Our approach offers a means for the precise and real-time control of visual features within an environment during conditioning – thereby providing quantitative control over an important contextual feature during learning. Such a methodology could also be used to examine the effects of within-session context changes. For example, future studies could examine the effects of within-session changes to the testing context. This methodology could also be extended for the presentation of unconditioned visual stimuli (e.g., birds of prey; c.f. De Franceschi et al., 2016).

It should be noted that a previous study failed to show that rats could discriminate between two contexts (one paired with foot shock and one without) when the only difference between the safe and unsafe contexts were visual features (Bucci, et al. 2002). This is in general agreement with other context discrimination studies, which demonstrate that rats have more difficulty discriminating between similar environments (few feature differences between context) than between explicitly different contexts; with enough training (e.g.,, > 14 days), however, rodents are eventually able to discriminate between two similar contexts (Nakashiba, et al. 2012). The discrepancy between our results and those of Bucci and colleagues (2002) may derive from differences in behavioral procedures. In the CPFE, context learning occurs in the absence of foot shock, whereas during discrimination learning context learning and foot shock association occur during the same session. We would predict that manipulations of visual features during context discrimination using our current design would initially replicate the findings reported by Bucci and colleagues (2002); no discrimination over the first 10 days), but with extended discrimination training (e.g., > 15 day; Nakashiba, et al. 2012), the rats would be able to discriminate between two contexts where the visual features predict the safe vs. unsafe context. However, more studies are needed to test these hypotheses.

Given the ease of implementation across laboratories of our preparation it is important to note some important caveats with an LCD based approach. First, we conducted our experiments using a Long Evans strain of rat. It will be important to see if our findings generalize to other rodents. For example, differences in fear conditioning (e.g., Chang & Maren, 2010) and innate freezing to predatory cues (Rosen, West, & Donley, 2006) have been reported across strains. Due to the visual component of our design, it will be necessary to determine if rodents with reduced visual acuity (e.g., Sprague-Dawley rats; (Prusky, Harker, Douglas, & Whishaw, 2002; Prusky, West, & Douglas, 2000) can detect different visual environments projected on LCD screens. Second, we found our effects with pumpkins and a rock formation (see figure 1). We have not tested other variations of visual images to exclude the possibility that our results are a selective phenomenon with these images or a general phenomenon with any images. However, other studies have successfully applied alternative methods to manipulate spatial navigation (Aghajan et al., 2014; Furtak et al., 2009; Harvey, Collman, Dombeck, & Tank, 2009), discrimination learning (Lee & Shin, 2012), and recently defensive behaviors (De Franceschi, Vivattanasarn, Saleem, & Solomon, 2016). Future studies should examine different visual images. Finally, we detected our effects by manipulating pre-exposure to the visual environment prior to immediate shock training in the CPFE paradigm – a paradigm which typically produces weaker conditioning under similar US parameters to single-trial contextual fear conditioning preparations (Schreiber, Asok, Jablonski, Rosen, & Stanton, 2014). Thus, it is unclear as to what degree rats would visually discriminate under single-trial CFC training and testing. An examination of context discrimination within our set-up (e.g., dCtx A+ [foot shock], dCtx B-[no foot shock]) is a logical next step to determine how post-conditioning manipulations of visual features of context affect behavior.

The present study offers a first step as a proof-of-concept in manipulating visual elements with LCD screens during incidental contextual learning in fear conditioning. More studies are needed to understand how rodents can visually discriminate between contexts in addition to how other elements (Rudy & O’Reilly, 1999) such as spatial (McHugh & Tonegawa, 2007), tactile, auditory, and olfactory (González, Quinn, & Fanselow, 2003) elements statistically contribute to contextual fear learning behaviorally and neurobiologically (Fanselow, 2010).

## Acknowledgements

This research was supported in part by the Oscar Kaplan Postdoctoral Fellowship in Developmental Issues awarded to Nathen J. Murawski. Additional support from 1 R01 HD075066-01A1 to ME Stanton and JB Rosen and the Department of Psychological and Brain Sciences, University of Delaware (AA). We thank Jamie Queensberry at the University of Delaware for help in fabricating the chamber. We also thank the University of Delaware Office of Laboratory Animal Medicine for care of the animals.

